# Stability of synchronous states in sparse neuronal networks

**DOI:** 10.1101/2020.02.02.931048

**Authors:** Afifurrahman, Ekkehard Ullner, Antonio Politi

## Abstract

The stability of synchronous states is analysed in the context of two populations of inhibitory and excitatory neurons, characterized by different pulse-widths. The problem is reduced to that of determining the eigenvalues of a suitable class of sparse random matrices, randomness being a consequence of the network structure. A detailed analysis, which includes also the study of finite-amplitude perturbations, is performed in the limit of narrow pulses, finding that the stability depends crucially on the relative pulse-width. This has implications for the overall property of the asynchronous (balanced) regime.

## I. INTRODUCTION

Networks of oscillators are widely studied in many fields: mechanical engineering [1, 2], power grids [3], arrays of Josephson junctions [4], cold atoms [5], neural networks [6], and so on. Such networks can be classified according to the single-unit dynamics, the coupling mechanism, the presence of heterogeneity, and network topology. Since phases are the most sensitive variables to any kind of perturbation [7], most of the attention is devoted to setups composed of phase oscillators [8], i.e. one-dimensional dynamical systems. However, even the study of such relatively simple models is not as staight-forward as it might appear. In fact, a wide variety of dynamical regimes can emerge even in mean field models of identical oscillators, ranging from full synchrony to splay states, and including hybrides, such as partial synchronisation [9], chimera [10], and cluster states [11]. General theory of synchronisation is, therefore, a much investigated field.

In this paper we focus on synchronous states by referring to a rather popular class of neural networks, but the whole formalism can be easily extended to more general systems so long as the coupling is mediated by the emission of pulses. In neuroscience the neuron dynamics is often described by a single variable, the membrane potential, which evolves according to a suitable velocity field. The resulting model is equivalent to a phase oscillator, where the variable of the bare system increases linearly in time while the complexity of the evolution rule is encoded in the phase response curve (PRC), which accounts for the mutual coupling [12]. Under the additional approximation of a weak coupling strength, the model can be further simplified and cast into a Kuramoto-Daido form, where the coupling depends on phase differences between pairs of oscillators [13, 14]. Here, however, we stick to pulse-coupled oscillators.

The stability of the synchronised state of pulse-coupled phase oscillators has been first studied in the context of excitatory *δ*-pulses [15]. Synchronisation is induced when two oscillators are sufficiently close and a common excitatory *δ*-pulse instantaneously sets both to the same value. Later, the stability analysis for excitatory and inhibitory pulses [16, 17] has been extended to *δ*-pulses with continuous PRCs [18]. General formulas are mostly available under severe restrictions, such as identical oscillators, mean field interactions, or *δ*-like pulses.

The *δ*-like pulse assumption is particularly limiting, not only because realistic systems are characterized by a finite width, but also because it has been shown that zero-pulsewidth is a singular limit, which does not commute with the thermodynamic limit (infinitely large networks) – at least in the context of splay states [19]. Relaxing the zero-width limit forces to increase the phase-space dimension to account for the dynamics of the fields felt by the different neurons. The most general result we are aware of is a formula derived in Ref. [20] for a single population of identical neurons in the presence of mean-field coupling and the so-called *α*-pulses.

The introduction of sparseness implies a significant increase in the computational complexity because of the randomness of the connections. In this context, the most relevant results are those derived in Ref. [21], where a sparse random network (Erdös-Rényi type) has been investigated in the presence of *δ*-pulses. The approach is rather complex since the noncommutativity associated with changes in the order of the incoming spikes obliged the authors to introduce a multitude of linear operators to solve the problem.

Here, we extend this kind of stability analysis to finite pulse-widths in two populations of excitatory, respectively inhibitory, neurons. Our approach can also be considered as an extension to sparse networks of the work in Ref. [20] devoted to mean-field models. This setup is chosen in studies of the so-called balanced state [22], where the asynchronous regime is dominated by strong fluctuations. Typically, the balance depends on both the relative size of the two populations and the relative amplitude of the pulses. In this paper, a careful study of the fully synchronous regime shows that also the relative pulse-width plays a non-trivial role.

Finite-width pulses can obviously have infinitely many different shapes. In this paper we consider the simplest case of exponential spikes and assume, as usual, that they superpose linearly. In practice, this means that each oscillator (neuron) is characterized by three variables: the phase or, equivalently, the membrane potential and two variables describing the incoming excitatory and inhibitory fields, respectively. At variance with Ref. [20], instead of transforming the model into a mapping (from one to the next spike emission), here we preserve the time continuity, as this approach allows for a more homogeneous treatment of the oscillators maintaining the full 3*N* dimensional structure of the phase-space (where *N* is the number of oscillators).

Furthermore, in agreement with previous publications [23–25] we assume that each neuron receives exactly the same number of excitatory and inhibitory synaptic connections. In fact, in spite of the random connectivity, in the fully synchronous regime, all neurons are characterized by exactly the same input. The degeneracy of the Lyapunov spectrum observed in mean-field models is lifted and the stability must be assessed by determining the eigenvalues of a suitable (sparse) random matrix.

More precisely, in Sec. II we define the model, including the specific phase response curve used to perform numerical tests. The overall linear stability analysis is discussed in Sec. III, first with reference to the general case and then specifically referring to short (but finite) pulses. In the same section we also determine the conditional Lyapunov exponent λ_*c*_, (i.e the exponent describing the response of a single neuron subject to a given - periodic - forcing): at variance with the mean-field model, λ_*c*_ differs from the maximum exponent of the whole network, indirectly confirming the nontrivial role played by the connectivity. In Sec. IV, we implement the formulas determined in the previous section to discuss the qualitative changes observed by varying the relative pulse-width. Finally, Sec. V is devoted to a summary of the main results and an outline of the open problems.

## II. MODEL

The object of study is a network of *N* phase-oscillators (also referred to as neurons), the first *N_e_* being excitatory, the last *N_i_* inhibitory (obviously, *N_e_* + *N_i_* = *N*). Each neuron is characterized by the phase-like variable Φ^*j*^ ≤ 1 (formally equivalent to the membrane potential), while the (directed) synaptic connections are represented by the connectivity matrix **G** with the entries

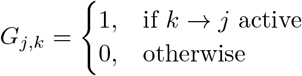

where 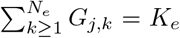 and 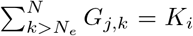, meaning that each neuron *j* is characterized by the same number of incoming excitatory and inhibitory connections, as customary assumed in the literature [23] (*K* = *K_e_* + *K_i_*, finally represents the connectivity altogether).

The evolution of the phase of both excitatory and inhibitory neurons is ruled by the same equation,

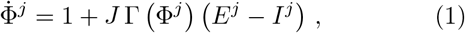

where Γ(Φ) represents the phase-response curve (PRC), *J* the coupling strength and *E^j^*(*I^j^*) the excitatory (in hibitory) field generated by the incoming connections. He we assume that *K* as well as *J* are independent of *N*, i.e. we refer to sparse networks. Whenever Φ^*j*^ reaches the threshold Φ_*th*_ = 1, the phase is reset to Φ_*r*_ = 0 and enters a refractory period *t_r_* during which it stands still and is insensitive to the action of the excitatory (*E^j^*) and inhibitory (*I^j^*) field. At the same time, the fields of the receiving neurons are activated. If the neuron *k*, emitting a spike at time 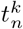, is excitatory (*k* ≤ *N_e_*), then the excitatory field *E^j^* of any receiving neuron *j* is activated (and similarly for the inhibitory field *I^j^*).

The fields in Eq. (1) evolve according to the differential equations

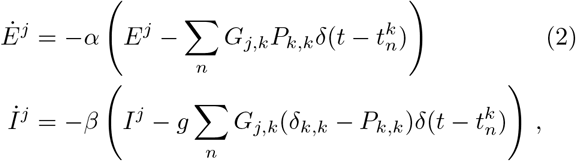

where *α*(*β*) denotes the inverse pulse-width of the excitatory (inhibitory) spikes. The coefficient *g* accounts for the relative amplitude of inhibitory spikes compared to excitatory ones. *P_k,m_* represents the elements of a projector operator **P**, separating excitatory from inhibitory neurons: *P_k,m_* = 0 except when *k* = *m* ≤ *N_e_*, in which case *P_k,k_* = 1.

In order to be more specific, we introduce the PRC used later on as a testbed for the formalism developed in the next section. We have chosen to work with the following piecewise linear PRC,

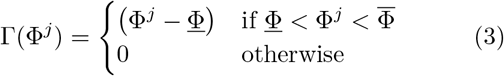

where <u>Φ</u> < 0, and 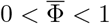 characterize the PRC. The resulting shape is plotted in Fig. 1 for <u>Φ</u> = −0.1 and 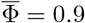 [26].

**FIG. 1.**
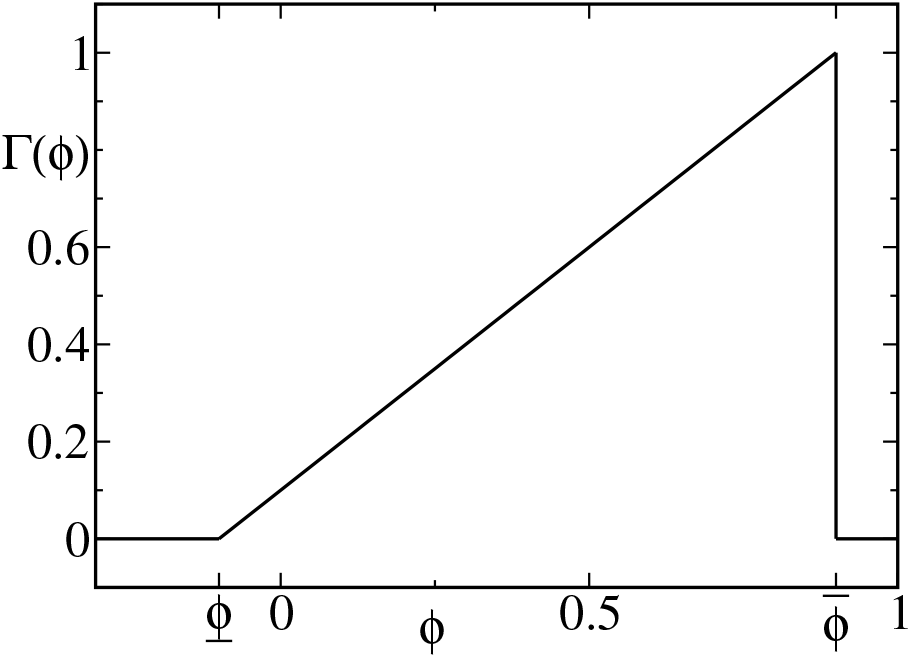
Example of the phase response curve (PRC) used in Sec. IV with 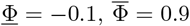 in combination with Φ_*r*_ = 0 and Φ_*th*_ = 1.

As anticipated in the introduction, we are interested in assessing the stability of the fully synchronous dynamics of period *T* as a function of the relative pulse-width, where *T* is the interspike interval. The solution is obtained by integrating the equation,

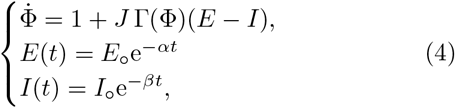

where

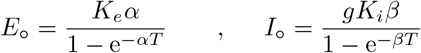

are the magnitudes of the fields immediately after the synchronous spike emission. The constants *E*_o_ and *I*_o_ result self-consistently from the sum of the remaining field at the end of the period *T* plus the contribution from the spike emission.

In the present paper, we focus on the stability of the synchronous period-1 solution (i.e. the initial configuration is exactly recovered after one spike emission). For long inhibitory pulses (small inhibitory decay rate *β*) we observed also stable period-2 and higher order periodic solutions. Fig. 2 shows the transition from stable period-1 to period-2 solution in the *α*, *β* plane (from top to bottom). Higher order periodic solutions appear underneath that curve in the shaded area and result in a synchronous bursting dynamics.

**FIG. 2.**
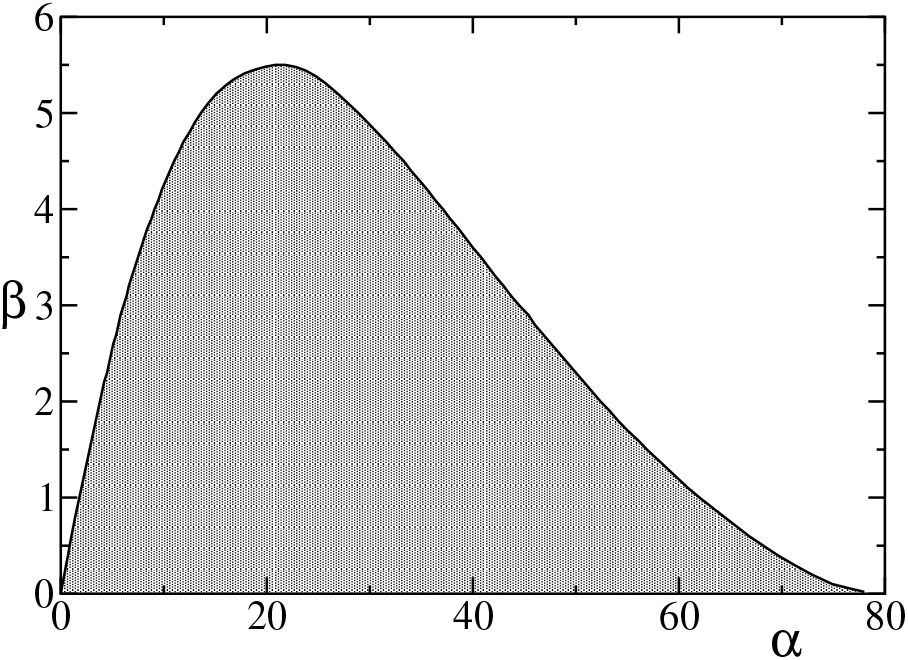
Phase diagram of synchronous regimes. Period-1 solutions exist above the curve in the white area. Crossing the curve, in the grey area, period-2 solutions first emerge followed (further down) by longer period regimes.

## III. LINEAR STABILITY ANALYSIS

### A. General theory

In this section we present the stability analysis of a synchronous state in the period-1 regime. At variance with Ref. [20], we do not construct the corresponding map, which means that the phase-space dimension is not reduced by a suitable Poincar’e section and the presence of a neutral direction is preserved. Actually this property can be even used to double check the correctness of the final result.

We start introducing a stroboscopic representation and focus on a weakly perturbed synchronous configuration

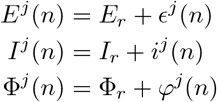

where all variables are determined at the end of consecutive refractory periods. As shown in Fig. 3 and clarified in the following, it is convenient to refer *φ^j^*(*n*) to one period later with respect to *ϵ^j^*(*n*) and *i^j^*(*n*). The fields *E_r_* = *E*_o_e^−*αt_r_*^, *I_r_* = *I*_o_e^−*βt_r_*^, and Φ_*r*_ = 0 do not depend on *n*, as the reference trajectory is periodic of period *T*. The overall perturbation can be represented as a 3*N* dimensional vector [***ϵ***(*n*), ***i***(*n*), ***φ***(*n*)]. For the future sake of simplicity, it is convenient to introduce also a second representation in terms of time shifts, ***v***(*n*) = [***τ***_*ϵ*_(*n*), ***τ***_*i*_(*n*), ***τ***_*φ*_(*n*)], where

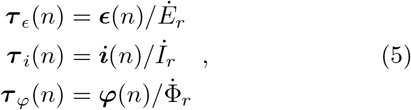

and 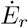, 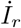 and 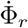 all denote time derivates at the end of a refractory period. In practice ***τ***_*x*_ corresponds to the time shift of the original trajectory to match the current perturbed state. The recursive transformation can be formally written as

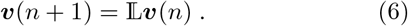

**FIG. 3.**
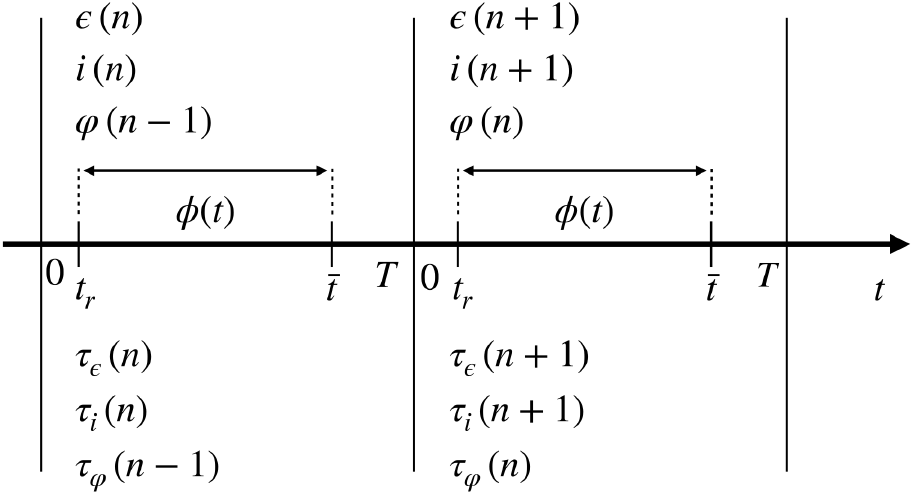
An illustration of the perturbation analysis in time *t* for the synchronous state.

Our next task is to determine the operator 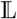. We start from the evolution equation of the excitatory field,

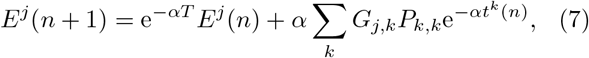

where *t^k^*(*n*) is the time elapsed since the arrival of the spike sent by the *k*th neuron in the *n*th iterate.

Since the trajectory is close to the synchronous periodic orbit, *E^j^*(*n* + 1) = *E_r_* + *ϵ^j^*(*n* + 1), and 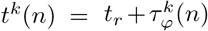. Up to first order in the perturbations, Eq. (7) yields,

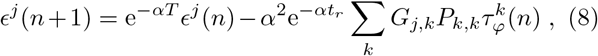

or, in vector notations,

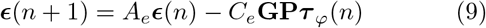

where

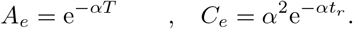

A similar analysis for the inhibitory field leads to

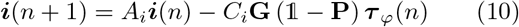

where 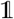 is the *N* × *N* identity matrix, while

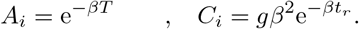

Notice that, at variance with the previous case, there is an extra factor *g* in the definition of *C_i_* to account for the larger amplitude of the inhibitory spikes.

Finally, we deal with phase dynamics. The core of the transformation is the mapping between the amplitude ***φ***(*n*) of the perturbation at time *t_r_* and the amplitude 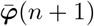 at time 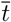, which can be formally written as

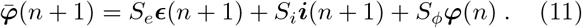

This transformation is diagonal (it is the same for all components); the three unknown parameters, *S_e_*, *S_i_*, and *S_ϕ_*, can be determined by integrating the equation obtained from the linearization of Eq. (1). To separate the notation of the stroboscopic phase perturbation *φ*(*n*) from the continuously developing phase perturbation between *t_r_* and 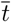 we introduce *ϕ*(*t*),

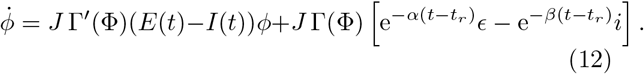

Upon setting [*ϵ,i,ϕ*(*t_r_*)] = [1, 0, 0], [0,1, 0], and [0, 0,1], 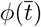 corresponds to *S_e_*, *S_i_*, and *S_ϕ_*, respectively. Once 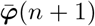 is known from Eq. (11), it can be transformed into the corresponding time shift

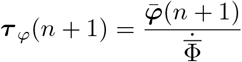

where

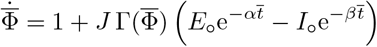

is the time derivative of the phase in the point where the neuron stops feeling the action of the field. In between 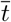 and *T*, the oscillators evolve with the same velocity and no adjustment of the time shift can be expected. The transformation is completed by Eq. (5), which allows mapping *φ*(*n*) onto the corresponding time shift and obtaining

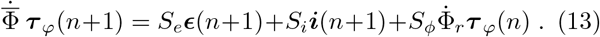

With the help of Eqs. (9,10), we find

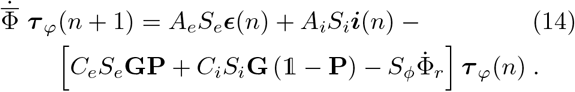

or, in a more compact form,

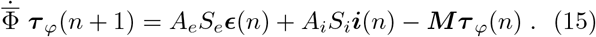

where **M** is an *N* × *N* matrix whose entries are defined as follows,

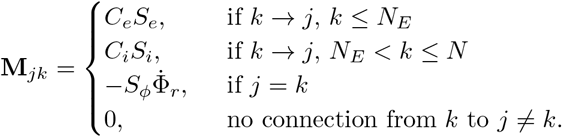

For homogeneity reasons, it is convenient to express all of the three recursive relations in terms of the components of the ***v*** vector,

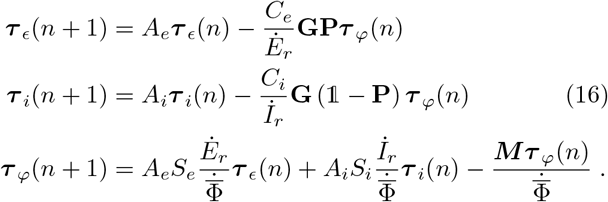

Now let us consider a homogeneous perturbation, such that ***τ***_*ϵ*_ = ***τ***_*i*_ = ***τ***_*φ*_. This perturbation must be mapped exactly onto itself, since it corresponds to a time shift of the whole orbit. Let us see what this amounts to. From the first of the above equations, we have that

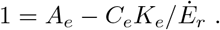

By looking at the definition of the various quantities, we can see that the equality is indeed satisfied. This is because *C_e_*/*Ė_r_* = −(1 − *A_e_*)/*K_e_*. Analogously, we can verify that *C_i_*/*İ_r_* = −(1 − *A_i_*)/*K_i_*, so that we can rewrite the transformation as

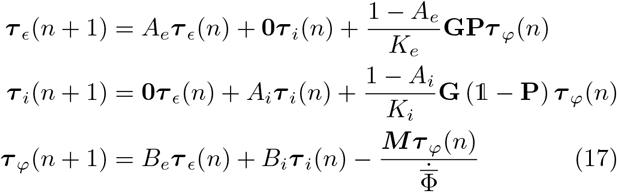

where 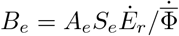 and 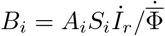.

By playing the same game of homogenous perturbations with the last equation of Eq. (16), we find that

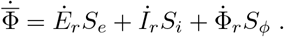

Direct numerical simulations confirm that this condition is satisfied, as it should, since it implies that a homogeneous shift of the phase of all oscillators is time invariant.

Altogether Eq. (17) is a representation of the linear operator 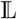 formally introduced in Eq. (6). The eigenvalues of 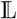 are the so-called Floquet multipliers *Z_i_*; the synchronous solution is stable if the modulus of all multipliers is smaller than 1[27]. One can equivalently refer to the Floquet exponents λ_*i*_ = log|*Z_i_*| that we also call Lyapunov exponents with a slight stretch of the notations.

For *α*, *β* ≫ 1 the fields are exponentially small when the neurons reach the threshold. In this limit, the fields behave as *slaved* variables and their contribution can be neglected in the stability analysis, which reduces to diagonalizing an *N* × *N* matrix,

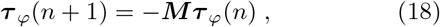

(notice that 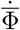 can be safely set equal to 1, as the coupling is negligible at time 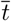).

### B. Transversal Lyapunov exponent

A simpler approach to assess the stability of the synchronous regime consists in investigating the stability of a single neuron subject to the external periodic modulation resulting from the network activity. The corresponding growth rate λ_*c*_ of infinitesimal perturbations is called transversal or conditional Lyapunov exponent. In mean-field models, this approach leads to the same result obtained by implementing a more rigorous theory which takes into account mutual coupling. Let the time shift at the end of a refractory period be equal to *τ_r_*; the corresponding phase shift is therefore

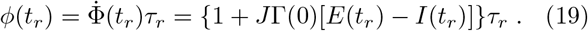

From time *t_r_* up to time 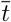 the phase shift evolves according to simplified version of Eq. (12),

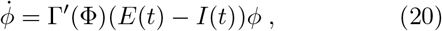

where we have neglected the variation of field dynamics, since the field is treated as an external forcing. As a result,

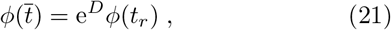

where, with reference to the PRC Eq. (3),

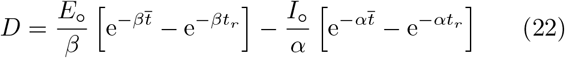

The corresponding time shift is

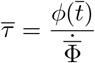

The shift 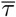 carries over unchanged until first the threshold *ϕ* = 1 is crossed and then the new refractory period ends. Accordingly, from Eqs. (19,21), the expansion *R* of the time shift over one period (a sort of Floquet multiplier) can be written as

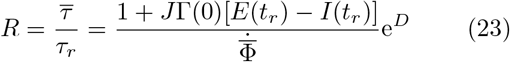

This formula is substantially equivalent to Eq. (54) of Ref. [20] (Λ_*ii*_ corresponds to *R*), obtained while studying a single population under the action of *α*-pulses. An additional marginal difference is that while in Ref. [20] the single neuron dynamics is described by a non uniform velocity field *F*(*x*) and homogeneous coupling strength, here we refer to a constant velocity and a phase-dependent PRC, Γ(*ϕ*).

The corresponding conditional Lyapunov exponent is

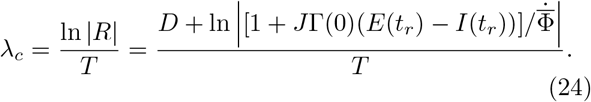

It is the sum of two contributions: the former one accounting for the linear stability of the phase evolution from reset to threshold (*D*/*T*); the latter term arises from the different velocity (frequency) exhibited at threshold and at the end of the refractory period. Notice the in the limit of short pulses, the field amplitude at time 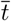 can be set equal to zero, thereby neglecting the corresponding exponential terms in Eq. (22) and assuming 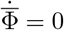.

## IV. APPLICATION

We now implement the general formalism in the case of the PRC defined by Eq. (3), considering a network with *N* = 1000 neurons, a 10% connectivity (i.e. *K* = 100 with *K_e_* = 80 and *K_i_* = 20), and *g* = 5; the coupling strength is assumed to be *J* = 0.03, while the refractory time is *t_r_* = 0.03. This setup, characterized by a slight prevalence of inhibition (*gK_i_* ≳ *K_e_*), is often adopted in the study of balanced regimes (see e.g. [23]).

The resulting Floquet spectra are presented in Fig. 4 for three different pairs of not-too-large *α* and *β* values. Rather than diagonalizing the matrix defined by Eq. (17), the 3,000 exponents have been determined by implementing a standard algorithm for the computation of Lyapunov exponents [28]. The larger are *α* and *β*, the more step-like is the spectral shape, the two lower steps being located around the decay rate (i.e. the inverse pulse-width) of the pulses (see the three horizontal dashed lines, which correspond to λ = −3, −4, and −8, respectively). This is sort of expected, since the field dynamics basically amounts to a relaxation process controlled by the respective decay rate. Anyhow, since the overall stability is determined by the largest exponents, it is sufficient to restrict the analysis to the first part of the spectrum (to the left of the vertical dashed line in Fig. 4), which, in the limit of large *α* and *β*, can be directly determined by diagonalizing the matrix defined in Eq. (18).

**FIG. 4.**
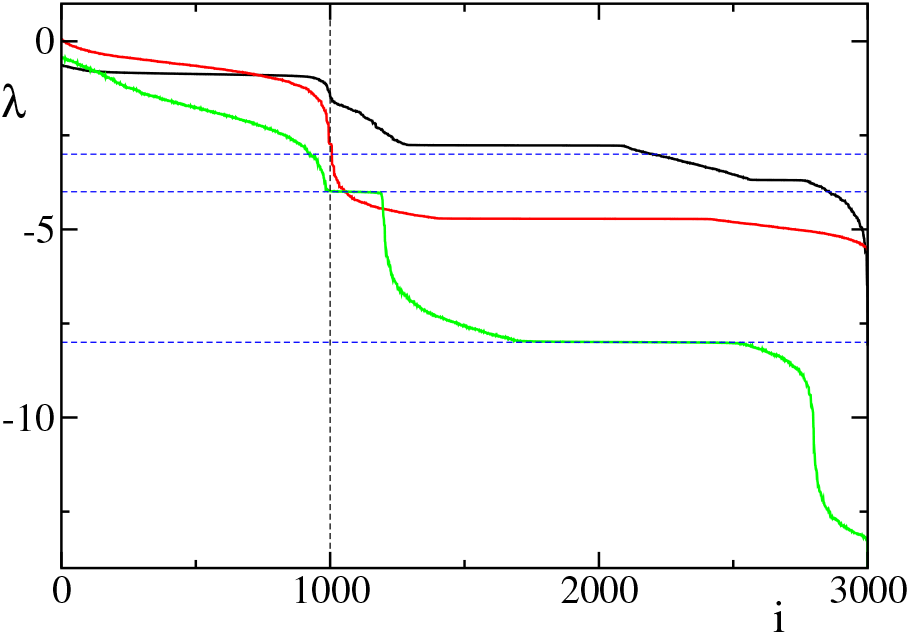
The total Lyapunov spectrum, for (*α*, *β*) equal to (4,3), (4,4), and (4,8) – top to bottom; the coupling strength is J = 0.03, as for all of our simulations, while *N* = 1000. The three horizontal dashed lines correspond to the three different rates used to identify *α* and *β* values. The vertical dashed line separates the part of the spectrum which, for large *α* and *β* values, is related to the actual network structure.

The dependence of the maximum exponent λ_*M*_ on the (inverse) pulse-width of the inhibitory spikes is reported in Fig. 5 (see the upper red curve). In this case, the Floquet exponent has been obtained by diagonalizing the matrix in Eq. (18) for a system size *N* = 10, 000 and a connectivity *K* = 1000 (*K_e_* = 800, *K_i_* = 200).

**FIG. 5.**
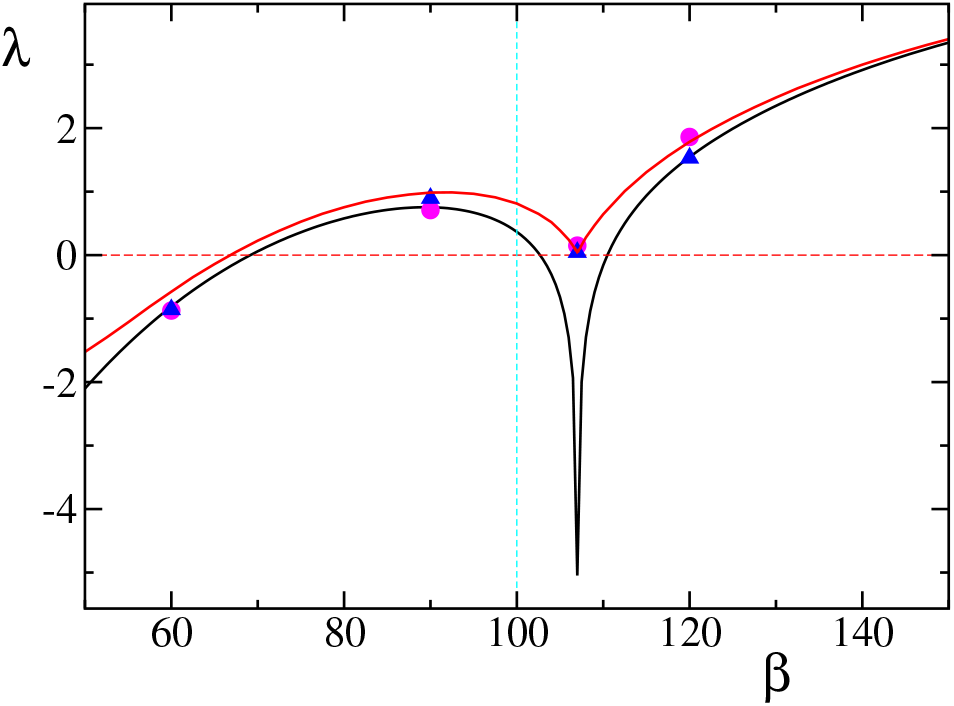
The maximal Lyapunov/Floquet exponent λ_*M*_ (upper red curve) vs. *β* for a network with *N* = 10, 000, *α* = 100 and the other parameters set as in the previous figure. The black curve corresponds to the transversal/conditional exponent λ_*c*_, while full dots and triangles result from the computation of the finite-amplitude Lyapunov exponent λ_*f*_ for *σ* = 10^−2^ and 10^−3^, respectively.

The vertical dashed line corresponds to the symmetric case, where both excitatory and inhibitory neurons have the same width (and shape). Interestingly, the stability, determined by the largest non zero exponent, (the always present λ = 0, corresponds to the neutral stability associated to a time shift of the trajectory) depends strongly on the relative excitatory/inhibitory pulse width and can even change sign: the synchronous solution is stable below *β* = 67 [29]. Additionally, there is evidence of a sort of singularity around *β* = 107, when the inhibitory spikes are slightly shorter than the excitatory ones.

Given the finite dimension of the matrices, sample-to-sample fluctuations are expected. Such fluctuations are, however, rather small, as testified by the smoothness of the red curve in Fig. 5. In fact, the single values of the Floquet exponents have been obtained not only by varying the (inhibitory) pulse-width, but also considering different network realizations. Although small, the fluctuations prevent drawing definite conclusions about the singularity seemingly displayed by the derivative of λ_*M*_(*β*) around *β* = 107.

In the limit of a fully connected network, we expect a perfectly degenerate spectrum (all directions are mutually equivalent) and λ_*M*_ equal to the conditional Lyapunov exponent λ_*c*_defined in Eq. (24). The lower black curve reported in Fig. 5 corresponds to λ_*c*_; except for a narrow region around *β* = 107, λ_*c*_ is always close to (lower than) λ_*M*_. This means that the mean-field approximation still works pretty well in a network of 10,000 neurons with a 10% connectivity.

The explicit formula Eq. (24) helps also to shed light on the *β* dependence of the network stability. The main responsible for the qualitative changes observed around *β* = 107 is the logarithmic term, arising from the difference between the velocity at threshold (equal to 1, irrespective of the *β*-value) and the velocity at the end of the refractory period. This latter velocity is determined by the effective field *E_eff_*(*t_r_*) = *E*(*t_r_*) – *I*(*t_r_*) which in turn strongly depends on the relative pulse-width. The time dependence of *E_eff_* can be appreciated in Fig. 6, where we report the trace for three different *β* values (60, 90, and 120) and the same *α* = 100. There, we see that even the sign of the effective field may change; for *β* = 120, Eeff is initially negative because inhibition dominates, but above *t* = 0.02 < *t_r_* the slower decay of the excitatory pulses takes over, so that the effective field amplitude is positive at the end of refractoriness. For *β* = 90 < *α* = 100, inhibition prevails at all times and the effective field is thereby negative for *t* = *t_r_*. Finally, for *β* = 60, excitation initially prevails, but inhibition takes soon over.

**FIG. 6.**
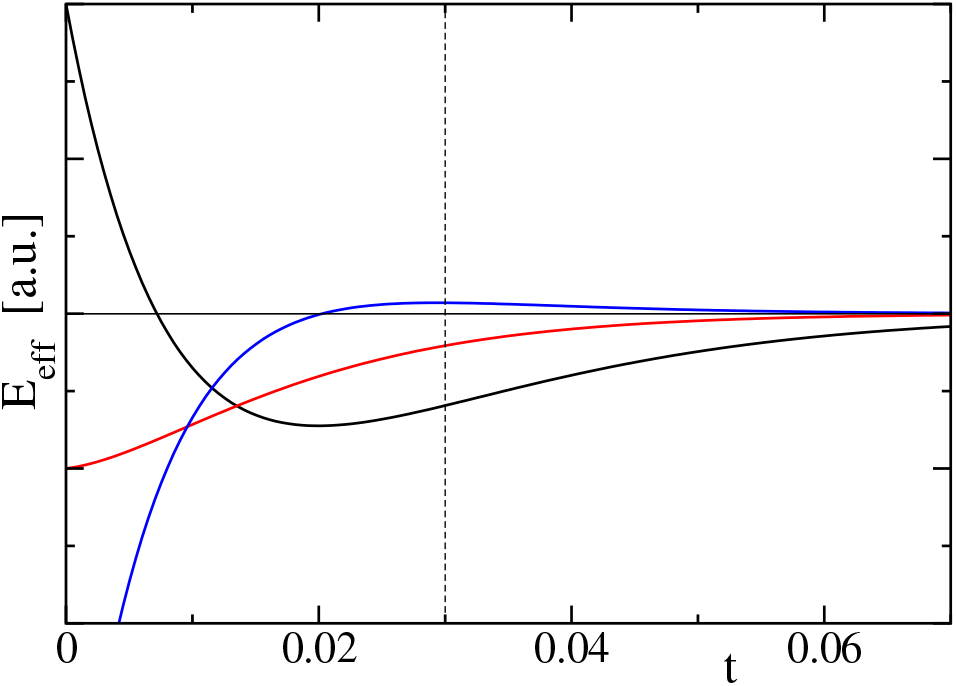
Effective field shape for *α* = 100 and *β* = 60 (black), *β* = 90 (red), and *β* = 120 (blue). The vertical dashed line identifies the end of the refractory period.

From Eq. (24), we see that the sign of the logarithmic contribution changes depending whether the argument is smaller or larger than 1. More precisely if the effective field is negative but larger than −2/(*J*Γ(0)), the discontinuity of the velocity tends to stabilize the synchronous regime; if *E_eff_*(*t_R_*) = −1/*J*Γ(0) the orbit is even superstable, i.e. the Lyapunov exponent is infinitely negative. This is precisely what happens for *β* ≈ 107. Altogether, the *β* interval around 107 separates the region where the expansion/contraction factor is positive (to the right), from the region where it is negative (to the left).

The sign of the multiplier has a simple explanation: 1 + *J*Γ(0) *E_eff_*(*t_r_*) < 0 means that the phase velocity is negative at the of the refractory period. Therefore, if one follows two nearby neurons – one leading over the other before reaching the threshold – then at the end of refractoriness, the leading neuron becomes the lagging one, as they initially move in the “wrong” direction[30]. This explains how the pulse-width may affect the stability.

So far we have referred to the Floquet exponents, without paying attention to the phase of the multipliers. In Fig. 7 we report both real and imaginary part of all eigenvalues for four different *β* values.

**FIG. 7.**
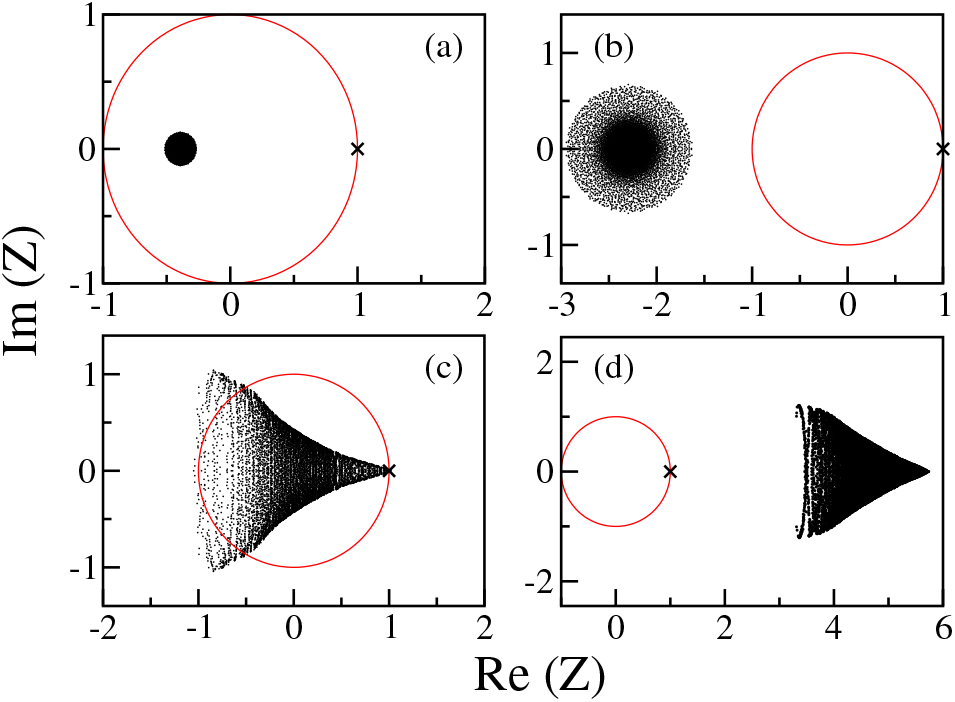
The distribution of the complex eigenvalues (black dots) for short pulses, *N* = 10,000, *α* = 100, and *J* = 0.03. The red curve highlights the unit circle. The black cross at (1, 0) singles out the always present eigenvalue associated with the periodic motion. The four panels represent the results for different decay rates of the inhibitory pulses *β*. In particular *β* = 60, 90, 107 and 120 for panel (a) - (d), respectively.

For *β* = 60 and 90, the eigenvalues (except for *Z* = (1, 0)) are distributed within a circle (see panels a and b). This is reminiscent of Girko’s theorem [31], which states that the eigenvalues of an *N* × *N* random matrix with independent and identically distributed entries (with zero mean zero and variance equal to 1/*N*), are uniformly distributed over the unit disc. However, it is not obvious how to adapt/extend this theorem to the present context, since the matrix **M** although being random does not satisfy several of the required assumptions, starting from the off-diagonal elements which take only three different values and their average is non zero.

Returning to Fig. 7, for *β* = 60 all the eigenvalues lie within the unit circle, meaning that the synchronous solution is stable, while for *β* = 90 *all* eigenvalues lie outside, meaning it is fully unstable: any perturbation is amplified!

Above *β* = 100, the spectrum changes shape, becoming funnel-like: for *β* = 120 (panel d), all eigenvalues sit again outside the unit circle, meaning that the synchronous solution is fully unstable. Interestingly, for *β* = 107 (panel c), the funnel is almost entirely contained inside the unit circle, so that the resulting (weak) instability is due to few complex eigenvalues lying on the upper-left and lower-left corners of the funnel. As an additional remark, we can see that the eigenvalue with largest modulus (i.e. the one determining the stability) is real and negative for *β* = 60 and 90, while it is real and positive for *β* = 120[32]. This is coherent with the behavior of the sign of the multiplier *R* (see Eq. (23)), which changes from positive to negative, while decreasing *β*. The qualitative differences observed in the region around *β* = 107 suggest that the “singular” behavior exhibited by λ_*M*_ is the signature of a true transition associated with a change of the spectral structure.

Finally, a few words about the leading eigenvector. It must possess some special features which are responsible for its larger expansion rate. However, we have not found any correlation with obvious indicators such as an anomalously large outgoing connectivity. We have only observed that the vector components are distributed in a Gaussian way with zero average.

### A. Finite-amplitude perturbations

Finally, we have directly investigated the stability of the synchronous regime, by studying the evolution of small but finite perturbations under the action of the model Eqs. (1–2) in the limit of short pulses. By following the same strategy developed in tangent space, the perturbation amplitude has been quantified as the tem-poral shift at a specific moment. We find it convenient to identify the *specific moment* with the threshold-passing time *t^L^*(*n*) of the last neuron (in the *n*th period). Provided the perturbation is small enough, all neurons are still in the refractory period and their phase is equal to 0 when the time is taken. The temporal shift of the *j*th neuron can be defined as *δ_j_* = *t^L^*(*n*) – *t_j_*(*n*), where *t_j_*(*n*) is its *n*th passing time. The perturbation amplitude is finally defined as the standard deviation *σ*(*n*) of all temporal shifts. Given an initial distribution with a fixed *σ*(0), it is let evolve to determine its value once the new set of spiking times is over. The ratio *R_f_* = *σ*(1)/*σ*(0) represents the contraction or expansion factor over one period *T*. Afterwards the standard deviation is rescaled to the original value *σ*(0) to avoid it becoming either too large to be affected by nonlinear effects or too small to be undetectable. We have found that *σ*(0) = 10^-3^, or 10^-2^ suffices to ensure meaningful results. The corresponding (finite amplitude) Lyapunov exponent λ_*f*_ is finally obtained by iterating this procedure to let the perturbation converge along the most expanding direction and thereby computing λ_*f*_ = ln|*R_f_*|/*T*. We have found that 50 iterates suffice to let the transient die out.

A crucial point is the integration time step, if the model is evolved by implementing an Euler algorithm. In fact, the time step must be much smaller than the separation between ocnsecutive spike-times, since they have to be well resolved. We have verified that setting the Euler integration time step Δ*t* at least 100 times smaller than *σ*(0) ensures a sufficient accuracy. The numerical results, plotted in Fig. 5 for four different *β* values (see the symbols), indeed confirm the theoretical predictions.

## V. CONCLUSIONS AND OPEN PROBLEMS

In this paper, we have developed a formalism to assess the stability of synchronous regimes in sparse networks of two populations of oscillators coupled via finite-width pulses. The problem is reduced to the determination of the spectral properties of a suitable class of sparse random matrices. Interestingly, we find that the relative width of excitatory and inhibitory spikes plays a crucial role even in the limit of narrow spikes, up to the point that the stability may qualitatively change. This confirms once more that the *δ*-spike limit is and it is necessary to include the spike width into the modelling of realistic neuronal networks.

Our analytical treatment has allowed constructing the stability matrix, but deriving an analytical solution of the spectral problem remains an open problem. The conditional Lyapunov exponent provides an approximate expression for the maximum Floquet exponent. It is quite accurate in a broad range of pulse-widths but fails to predict the weak instability occurring when inhibitory pulses are slightly narrower than excitatory ones. For relatively wider inhibitory pulses, numerical simulations suggest that it will be worth exploring the possibility to extend the circular law of random matrices to sparse matrices of the type herewith derived.

While mean-field models are characterized by a degenerate spectrum (all directions being equally stable), here the degeneracy is lifted by the randomness associated with the sparse connectivity. It is therefore desirable to understand which features make some directions so special as to be characterized by a minimal stability. This is probably related to the presence of closed loops of connections among oscillators which sustain an anticipated or retarded firing activity. Further studies are required.

## ACKNOWLEDGMENTS

Afifurrahman was supported by an LPDP Indonesia fellowship.

